# Assessment of self stigma and medication adherence among patients with mental illness who had been treated at Jimma University Specialized Hospital, South west Ethiopia: community based cross sectional study, 2017

**DOI:** 10.1101/340588

**Authors:** Eba Abdisa Golja, Shimelis Girma

**Author notes:** Corresponding author: lecturer and clinician at Wollega University, Nekemte, Ethiopia.

## Abstract

**Background:** self Stigma associated with psychiatric disorders and its treatment toward patients with mental illness is the main challengers for early and successful treatment outcome.

**Objective:** To assess the association between self stigma and level of medication adherence among patients with mental illness who have been treated at psychiatric clinic, JUSH, South West Ethiopia, 2017.

**Methods:** This study is a community based cross sectional study design using a case tracing method to identify patients in the community. To get the study subject, tracing method was carried to get the patients who had history of treatment at psychiatric clinic, Jimma University Teaching Hospital. Interview based Questionnaire was used to get information from the participants. The relationship was checked by doing both bivariate and multi variables linear regression.

**Result:** Of the total respondents, males were185 (61.7%), females 115 (38.3%), and the mean age of the respondents was 34.99 (SD=11.51) years. From the total (n=300) respondents, more than half of them, 182 (60.7%) were adherent to their psychotropic medication. Among adherent respondents, of all types of stigma, minimal stigma was 63.2% whereas moderate to high stigma was 22.4%.

**Conclusion:** It was identified that there was high self stigma related to medication non adherent among patients with mental illness. Therefore, mental health professionals and the health care policy shall work to increase the level of psychotropic medication adherence so as to decrease self stigma associated with mental illness.

## Introduction

Internalized Stigma and perceived discrimination associated with psychiatric illness and its treatment among people with mental illnesses which frequently results from them are main obstacles preventing early and successful recovery [1]. Stigma of psychiatric patient is a term that applies to labeling patients as different and inferior while discrimination is a behavioral manifestation of stigma and it is omission of patients from certain competitive areas, working as a form of intangible control over groups of people with mental illness [2]. In a particular, Internalized stigma refers to the process by which individuals with mental illness apply negative stereotypes to themselves, expect to be rejected by others, and feel alienated from society [3].

Self stigma and discrimination associated with mental illnesses have remained a global public health concern over the years [4]. Majority of studies on internalized stigma among individuals with mental illness originated from developed nations showed that the prevalence ranges from 20%-30% [5]. Brohan et al, in 2004, assessed the levels of self stigma among patients with mood disorders in 13 countries and they found that moderate to high levels of internalized stigma were present in 21.7% of individuals [6]. Ritsher and Phelan found that in USA the prevalence of elevated internalized stigma among severely mentally ill outpatients was 28% [7].

Studies from developing nations showed that around a quarter of patients with mental illness experience internalized stigma [8]. A study from Nigeria among outpatient psychiatric patients reported a prevalence of 21.6% [9]. Twenty five percent of Taiwanese patients with depressive disorders had experienced internalized stigma [10]. A study from Bangalore among schizophrenic patients showed that 93 percent of the individuals experienced stigma during their acute phase of illness [11]. A study from Mumbai reported that 69% of patients with schizophrenia experienced stigma in their personal lives [12]. A retrospective study conducted in Hyderabad in an out-patient psychiatric department of a University hospital showed that internalized stigma was present among 47% of individuals with various mental disorders [13]. Therefore, stigmatization and discrimination against persons with mental illness is a matter of public health concern globally. This is due to the fact that many people with mental illness are challenged twice; on one hand, they struggle with the symptoms and disabilities that result from the disease and on the other, they are stereotyped and prejudiced due to misconceptions about mental illness [14]. It is estimated that about one million people die due to suicide every year and one in four people worldwide will experience mental illness in her or his lifetime and is associated with self stigma and discrimination [15].

Results from epidemiologic studies suggest that 50–60% of individuals who could benefit from treatment do not seek care and leaving millions of individuals in service gap [16]. Perhaps even more concerning, an estimated 20% of children and adolescents struggle with a mental illness, yet 70% fail to receive adequate treatment [17]. Many factors contribute to this failure. We focus on one prevailing theory – that stigma toward mental illness may act as a significant barrier to seeking mental health care for both adults and adolescents [18].

According to the World Health Organization, the self stigma and discrimination that exist towards people with mental health problems is one of the most important issues to overcome in mental health policy [19]. There is no known country, society or culture in which people with mental illness are considered to have the same value and to be as acceptable as people who do not have mental illness [20].

As a consequence of stigma, patient with mental illness usually can have difficulty in maintaining their day-to-day social interactions, which in the worst case may result in them committing suicide [21]. Internalized stigma is not only a consequence of mental illness but also a factor that interferes with help-seeking behavior, and it may delay treatment-seeking in patients with mental illness with a lots said that the stigma of mental illness is more disabling than the illness itself [22]. It is a barrier for mentally ill patients to avail treatment early. As a result, they prolong their recovery period, experience complications, and face serious financial difficulties [23].

On the other hands, discrimination affects all areas of living. It can prevent people from getting or having basic things that most of us take for granted, like: Getting hired, promoted or keeping a job, Finding or keeping a place to live in a safe, accepting community, Getting proper health care, Feeling loved, needed and accepted by family and friends, Contributing to communities and feeling productive, Getting insurance, Going to school, Immigrating to another country, Feeling positive about ourselves [24].

In recent decades, there have been considerable developments in mental health, however, majority of the patients don’t seek treatment because of the stigma and discrimination [5]. Finding show that 50–60% of adult individuals who could benefit from treatment doesn’t seek care [18]. An estimated 20% of children and adolescents struggle with a mental illness, yet 70% fail to receive adequate treatment [19].

Lack of treatment means, either they never visited health setting for help or they may be non adherent to their prescribed medication. According to Adewuya, et al, the reason for medication non adherence among psychiatric patients could be a number of factors such as: discomfort resulting from treatment (side effects of medication), expense of treatment, problems of filling a prescription, decision based on personal value, religious or cultural beliefs, maladaptive personality, coping style, stigma and discrimination [12].

Researchers indicated that self stigma might be a potentially important factor contributing to non-adherence to medication. Which, in turn, non-adherence leads to increased hospitalization, higher healthcare costs and is a predictor of poor outcome related to relapse rates, progressive brain damage, suicide and overall mortality [25].A clinical assumption is that both patient experienced discrimination and anticipated stigma increase non-adherence since negative attitudes, shame and social withdrawal could cause failure to access healthcare. Self stigma was identified as an important barrier to adherence in some qualitative study [26].

Fear of stigma connected to the diagnosis and being prescribed with medication and fear of rejection due to revealing symptoms have been suggested to further increase non-adherence to psychiatric help [27]. In a qualitative study patients reported that the stigma of taking medication caused them to hide their medication from others and to take it only in privacy because of fear of social and self stigma [28].

Different studies showed that to reduce level of internalized stigma and perceived discrimination, different measures like training, education, contact, increasing health service coverage, mass media campaigns and a broad range of multifaceted interventions [29].

Many researchers also tried to dig magnitude of any types of stigmas and discriminations among mentally ill patients. There are also some findings that give emphasis on self stigma and discrimination as one of predictors of psychiatric treatment and to be compliant to their prescribed treatment but few was known about relation of level of self stigma and discrimination with level medication adherence.

Therefore, the aim of this study was to examine the association of self stigma with that of level of medication adherence among patients with mental illness treated at psychiatric clinic.

This study is important because it will come with prevalence of internalized stigma and discrimination among patients with mental illness who had history of treatment at psychiatric clinic, JUSH. It will inform new information on effect of psychiatric treatment on level of self stigma and discrimination. The result of the study will help the mental health workers, psychiatrists and others coworkers to give focus not only on signs and symptoms of the problem but also on self stigma and discrimination associated with mental illness.

The study will have great importance to the health sectors of the area, health bureau and officials to provide better services on reduction and prevention of internalized stigma and perceived discrimination which takes places on patients with mental illness. It will provide information (magnitude of self stigma and discrimination, effect of medication adherence on level of self stigma and discrimination) for the future further research on self stigma and discrimination related to medication adherence level in the area, in particular, and others part of the country, in general.

## Methods and materials

The study was conducted from April 20 to June24, 2017 at community level in Jimma town on individuals with mental illness. Jimma town is capital city of Jimma zone which was founded in the late 1830’s and it occurs at south west part of Ethiopia at a distance of 352 Km south west from Addis Ababa. Jimma Town is located 352 km southwest of Addis Ababa. Psychiatric outpatient and inpatient services are available only at the University hospital. From this town, there are more than 1200 individuals who have ever follow-up treatment at psychiatric clinics with different diagnosis and different duration of treatment. Out of this, 698 patients were adult patients.

### Study design

A community based cross sectional study design was applied.

### Eligible

Those adult patients who ever started psychiatric treatment at least once at psychiatric clinic, Jimma University teaching hospital

### Sample size determination

The actual sample size of the study will be determined using the formula of single population proportion formula.

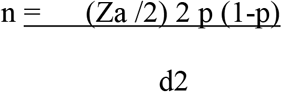

Where n = estimated sample size

Za/2 = Critical value at 95% confidence level of certainty (1.96)

P = population proportion

d = marginal error

To determine the sample size the following assumption were used. The previous finding reported that in near to this area the self stigma was 25.12%, so that we took the population proportion is 25.12% for our research. Therefore, the sample size by the above calculation is 289. By adding 10 % of non response rate, the final sample size is 318. But 14 of them were died from different physical illness which was the sequella of their mental illness. Therefore, we excluded them from the participants and the analysis was carried out among 304 participants who were traced and alive during the data collection whether they were interested or refused the participation.

### Sampling method and technique

To get study participants, simple random method was used because the registration of the patients with mental illness who are only from the town was used for different information. The total adult patients with mental illness were 1200 and from this total we selected 318 participants by using computer generated method. By reviewing chart of the patients, some recorded information (like: name, age, sex, address (Kebele), year of first visit, types of psychiatric diagnosis, etc) were identified. Finally, the information of adult psychiatric patients was distributed to health extension worker of the town and psychiatry department staffs according to the patients’ address. Health extension workers and psychiatry department staff identified patients whether they were dead, alive or change the place of residence by using tracing method as well as by using phone number of the patients which registered on the chart.

### Dependant variables

Self stigma

### Independent variables

Primary exposure variables

Level of medication adherence

Other predicting factors

➢ Sociodemographic characteristics
➢ Types of psychiatric diagnosis

Bipolar disorder
✤ major depressive disorder
✤ schizophrenia
✤ brief psychotic disorder
✤ Schizophreniform disorder
✤ schizoaffective disorder
✤ Others ……………
✤ delusional disorder
✤ panic disorder
✤ generalized anxiety disorder
✤ post traumatic stress disorder
✤ acute stress disorder
✤ social phobia
✓ Duration of treatments, premorbid occupational status, status of functional outcome
✓ Duration of the illness, age at onset of the illness,
✓ Number of episode, number of admission, Level of medication adherence, Social support

### Data collection tools

The data was collected using interviewer-administered questionnaires.

The questionnaire has five components: background characteristics, clinical inventory part, Internalized stigma of mental illness (ISMI), WHODAS version 2 (12 item), Oslo scale for social support and Morisky Medication Adherence Scales (MMAS-4)

Clinical inventory part include patients’ working diagnosis, age at onset, duration of illness, no of episode, no of hospitalization, duration of treatment, current clinical status, and previous suicide attempt. To measure PWMI self-stigma, the29-item Likert scale of Internalized Stigma of Mental Illness (ISMI) Scale will be used. In each measure, the items will be summed to get total scores of stigma so that higher scores indicated higher stigmas. The ISMI scale have a total of 29 items on a 4-point Likert (1 = strongly agree to 4 = strongly disagree) measure containing five subscales; Alienation (6 items), Stereotype Endorsement (7 items), Discrimination Experience (5 items), Social Withdrawal (6 items), and Stigma Resistance (5 items). Alienation is “the subjective experience of being less than a full member of society”. The Stereotype Endorsement is “the degree to which patients agreed with common stereotypes about people with a mental illness”. The Discrimination Experience measures “respondents’ perceptions of the way they tend to be treated by others”. The Social Withdrawal measures the self exclusion from social events/situation due to mental illness”. The Stigma Resistance subscale is “a person’s ability to resist stigma” and the higher score indicate the lower resistance of stigma [31].

The other tool which was developed by the World Health Organization (WHODAS-II) was used to establish level of impairment associated with mental illness and it assesses the level of disability and the number of days lost from work in the previous 30 days. It has 12 items and Likert scale (1-5). Oslo scale for social support has three items and is used to assess level of social support of the participants [32]. Morisky Medication Adherence Scales has four items and each of the item has dichotomous types of response (yes, no); the cutoff point of the scale is two; less than or equal to one indicate that adherent and more than or equal two indicates non adherent and was reliable in the study (Cronbach’s α=0.73) done in Jimma university specialized [30, 31].

### Data collection procedure

Data was collected by using interview based structured questionnaires among the participants those were addressed by tracing method in each kebele of the town. Two psychiatric nurses and four mental health masters recruited in data collection. Two instructors from psychiatry department were recruited in each kebele of the town as supervisor with principal investigator.

### Data quality control

To ensure quality of data, data collection tools which was used to assess outcome variables was that used previously in the same study area. Data collectors and supervisors were trained for two days on the purpose of the study, details of the questionnaires, technique of interview, importance of privacy and confidentiality of the respondents. The data collection instrument tools were developed in English version, then translated to Amharic and later translated back to English version by others experts to see its consistency. Twenty patients were pretested at psychiatric clinic, Jimma University teaching Hospital, those who came for appointment from different part of Jimma Zone except Jimma Town. During data collection procedures, all the collected data was reviewed and checked daily for its completeness.

### Data analysis and interpretations

The data was checked, coded and entered into Epi-data version 3.1, and then was exported to SPSS window version 21 for summarization and further analysis. Prior to analysis linear regression assumptions were checked. Simple linear regression analysis was carried out to assess association between each dependent and independent variables and to identify candidates for multivariate linear regression analysis. Variable having p-value less than 0.25 was subjected for multivariate linear regression analysis. Then, multivariate linear regression analysis was performed to get the final model. Statistical significance was considered at p-values less than 0.05 and adjusted odd ratio of 95% confidence interval.

A frequency table was computed for socio-demographic and other variables. Self stigma and perceived discrimination was identified against medication adherence status. Mean score for each of the subscale and overall self stigma was done but the final subscale of ISMI was reversely recoded before identified mean score. Finally, the result was presented in figure, tables, graphs and statements

### Ethical clearance

The study was approved and ethical clearance letter was obtained from institutional review board of college of health sciences, Jimma University and was given to Jimma town administrative office to undertake formal investigations. A consent sheet was prepared and translated to local language and attached to the questionnaire in a separate page. In the consent sheet, the purpose of this study was clearly stated and there was explanations that there was no way to cause any harm to study subjects and it indicated that there was no chance the participants’ identifiers to be written on the survey questionnaire and that no individual response would be reported. Written consent was obtained from study participants to ensure confidentiality. Participation was according to voluntarily of participants. For those individuals who seriously ill due to stopping their treatment, we connected them to psychiatric clinic to resume their treatment and others health care service by giving psycho education for both family and the patients.

### Dissemination plan

The result was presented to Jimma University; college of health sciences, department of psychiatry and documents was disseminated to all responsible bodies in the study area. The thesis was disseminated in to the ministry of health, Oromia health Bureau, Jimma zone health Bureau, Jimma University department of psychiatry. Hard and soft copy will be availed in the library of Jimma University for graduate students as well as for others concerned bodies to review when in need. The finding was presented at various seminars, workshops. Effort will be made to publish on national or international peer reviewed scientific journal.

### Operational definition

Level of stigma is based on the mean score of ISMI, and using similar score categories to the European and Ethiopian study [51]:

➢ <2 of total score indicated that minimal internalized stigma.
➢ 2-2.5 of total score indicated that low internalized stigma
➢ 2.5-3 of total score indicated that moderate internalized stigma
➢ 3+ of total score indicated that high internalized stigma
➢ Adherence to a medication is the extent to which patients take medications as prescribed by their health care providers. The level of drug adherence is measured based on MMAS score [52]. Accordingly, Adherence-MMAS score of ≤ 1, Non-adherence-MMAS scores of ≥ 2

## Results

### Background characteristics of the respondents

The total participants addressed by tracers were 318. From theses 14 people were died from physical illnesses and only 304 participants were included in the study. The response rate was 98.67% in this study. Of the total respondents, males were185 (61.7%), females 115 (38.3%), mean age of the respondents was 34.99 (SD=11.51) years. One hundred and eighty five (61.0%) were single. Majority of the study respondents were Muslim, which was 129 (43.0%) of all. One hundred forty five (48.3%) of them were Oromo, and 192 (64.2%) of them were living with their family. One hundred forty (46.7%) of the respondents were jobless and 54 (18.0%) were employed either in governmental or private organizations. Education level of respondents was primary level, 105 (35.0%), followed by secondary level which was 87 (29.0%), but those who followed their college and above education were only 83 (27.7%) of the total respondents. Average monthly income of respondents was 503.42 (SD=958.15) with minimum of zero to five thousand birr, but majority of the respondents [(228 (76.0%)] were earning monthly less than 500 ETB. Mean value of number of family members living with respondents was 4.48 (SD=2.197) with minimum of living alone to the highest of living 12 numbers of family members.

### Clinical characteristics of the respondents

Of the total respondents, one third (32%) them had working diagnosis of schizophrenia followed by major depressive disorder (24.3%). Average value of the duration of respondents experienced mental illness was 98.25 (SD=67.608) months with the minimum duration of 3 months to the maximum of 336 months. Mean value of the time duration that the respondents were staying without searching for treatment was14.28 (SD=24.31) months with range of visiting health setting immediately of illness to waiting 192 months without visiting. Average of time duration that respondents had been getting treatment was 69.11 (SD=64.977) months, with the minimum of zero month to the 348 months duration range. After the first episode of their illness, respondents experienced average value of 2.17 (SD=2.511) times relapse of their illness with the maximum value of 17 times. Number of admission respondents to hospital with average value of 0.87 (SD=1.289) within the range of no admission which was 54.0% to the value of seven times admission. Mean age of the participants before the onset of mental illness was 26.90 with standard deviation of 11.484 years.

Around 75% 0f the respondents were on the work (having jobs) when they faced their first illness, which were 224 (74.7%) and around one third of them were married. More than half of the respondents had history of discontinuation of their medication without medical advice which was 63.3%. Fifty five (18.3%) respondents had poor social supports whereas 39 (13.0%) of them had high social supports. Of all respondents, 30.7% attempt suicide with very short period of time with no strong intention to kill themselves to the extent of multiple attempts with strong intent. Forty seven percent of respondents attempted suicide due their illness alone whereas 44.8% of them attempted due to their illness comorbid with social problems. More than two third (69.3%) of the respondents had current follow up from psychiatric clinic, but those who were on treatment were 63.3% from them.

### Self Stigma among people with mental illness

Overall ISMI scale contains 29-items with four point Likert scale and has five subscales. The mean score of each subscale was done and accordingly, mean alienation (being inferior and contains five components) score of 2.16 with standard deviation of 0.95, stereotype endorsement contains seven components with four point scale and the mean score was 2.11 with standard deviation of 0.844, perceived discrimination contains five components with four point scale and the mean was 2.16 with standard deviation of 0.90, social withdrawal Contains six components with four point scale and mean was 2.10 with standard deviation of 0.857, stigma resistance contains five components with four point scale and mean was 2.11 with standard deviation of 0.844.

Overall ISMI contains five components (alienation, stereotype endorsement, perceived discrimination, social withdrawal and stigma resistance) with a 29 items; each has the four point scale. Stigma resistance component was reversely recoded because the maximum value indicates that resistance is low in opposite of others components of IMSI. In this finding, the mean value of overall self stigma was 2.15 with standard deviation of 0.707 and around one third (28%) of the respondents had moderate to high self stigma.

### Level of medication adherence

Medication adherence was assessed using Moresky scale which contains four components. It assessed level of medication adherence among individuals those who had been on treatment. From the total (n=300) respondents, more than half of them, 182 (60.7%) were adherent to their psychotropic medication. Those individuals forgetting taking of their medication at right time and right dose were 42.3% but those who stopped their medication when their illness become worse were only 17.3% and from them, 26% of the non adherents were males whereas females were 13.3%. Tables4. Summarize the number and percentage of Morisky components found in this research.

**Table 1.** Background characteristics of people with mental illness living in Jimma Town and have been treated at psychiatric clinic, JUSH, Southwest Ethiopia, 2017.

| Characteristic |  | Frequency (n=) | Percent (%=) |
| --- | --- | --- | --- |
| Sex | Male | 185 | 61.7 |
|  | Female | 115 | 38.3 |
| Marital status | single | 180 | 60.7 |
|  | married | 80 | 26.7 |
|  | divorced | 22 | 7.3 |
|  | widowed | 11 | 3.7 |
|  | separated | 5 | 1.7 |
| Religion | Orthodox | 121 | 40.3 |
|  | Muslim | 129 | 43.0 |
|  | Protestant | 44 | 14.7 |
|  | Catholic | 4 | 1.3 |
|  | Others<sup>*a</sup> | 2 | 0.7 |
| ethnicity | Oromo | 145 | 48.3 |
|  | Amhara | 75 | 25.0 |
|  | Dawuro | 28 | 9.3 |
|  | Gurage | 21 | 7.0 |
|  | Other<sup>*b</sup> | 31 | 10.3 |
| Living condition | with family | 194 | 64.7 |
|  | living alone | 17 | 5.7 |
|  | living on street | 5 | 1.7 |
|  | with relatives/friends | 8 | 2.7 |
|  | with wife and kids | 58 | 19.4 |
|  | other<sup>*c</sup> | 18 | 6.0 |
| Occupation | jobless | 139 | 46.3 |
|  | employed | 54 | 18.0 |
|  | retired | 7 | 2.3 |
|  | merchant | 26 | 8.7 |
|  | student daily | 29 | 9.7 |
|  | labor worker | 16 | 5.3 |
|  | house wife | 23 | 7.7 |
|  | other<sup>*d</sup> | 6 | 2.0 |
| Educational status | Illiterates | 19 | 6.3 |
|  | able to read/write, but no formal education | 6 | 2.0 |
|  | primary level | 103 | 34.3 |
|  | secondary level | 89 | 29.7 |
|  | college and above | 83 | 27.7 |

<sup>\*a</sup>adventist, catholic, Wakeffata <sup>\*b</sup>Yem, Kefa, Sidamo, Wolayita <sup>\*c</sup>in prison, dormitory <sup>\*d</sup>farmer

**Table2.**
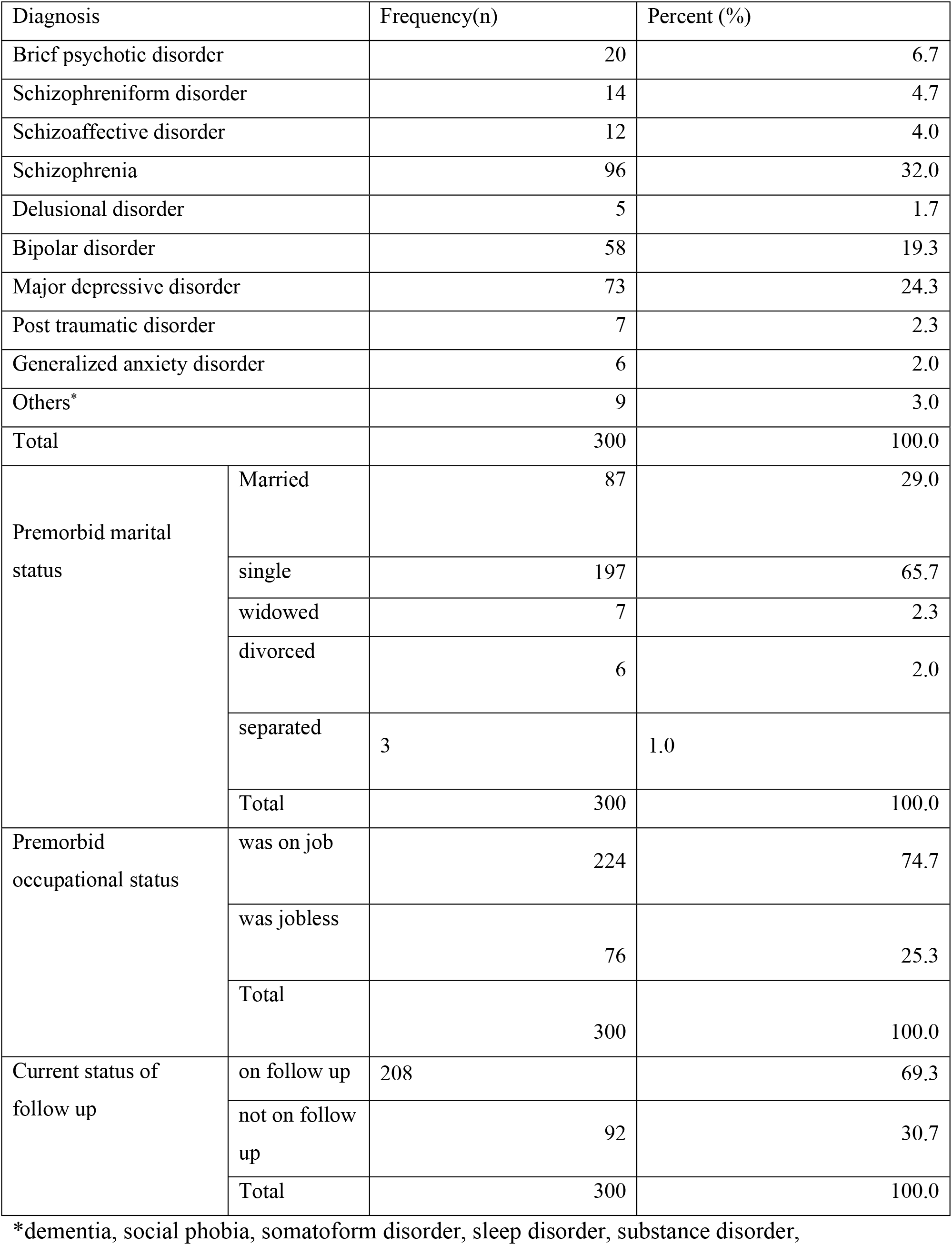
Working diagnosis and Clinical related characteristics of the patients with mental illness who had treatment at psychiatric clinic, 2017.

| Diagnosis | Frequency(n) | Percent (%) |
| --- | --- | --- |

|  |  |  |  |
| --- | --- | --- | --- |
| Brief psychotic disorder |  | 20 | 6.7 |
| Schizophreniform disorder |  | 14 | 4.7 |
| Schizoaffective disorder |  | 12 | 4.0 |
| Schizophrenia |  | 96 | 32.0 |
| Delusional disorder |  | 5 | 1.7 |
| Bipolar disorder |  | 58 | 19.3 |
| Major depressive disorder |  | 73 | 24.3 |
| Post traumatic disorder |  | 7 | 2.3 |
| Generalized anxiety disorder |  | 6 | 2.0 |
| Others* |  | 9 | 3.0 |
| Total |  | 300 | 100.0 |
| Premorbid marital status | Married | 87 | 29.0 |
|  | single | 197 | 65.7 |
|  | widowed | 7 | 2.3 |
|  | divorced | 6 | 2.0 |
|  | separated | 3 | 1.0 |
|  | Total | 300 | 100.0 |
| Premorbid occupational status | was on job | 224 | 74.7 |
|  | was jobless | 76 | 25.3 |
|  | Total | 300 | 100.0 |
| Current status of follow up | on follow up | 208 | 69.3 |
|  | not on follow up | 92 | 30.7 |
|  | Total | 300 | 100.0 |

\*dementia, social phobia, somatoform disorder, sleep disorder, substance disorder,

**Table3.**
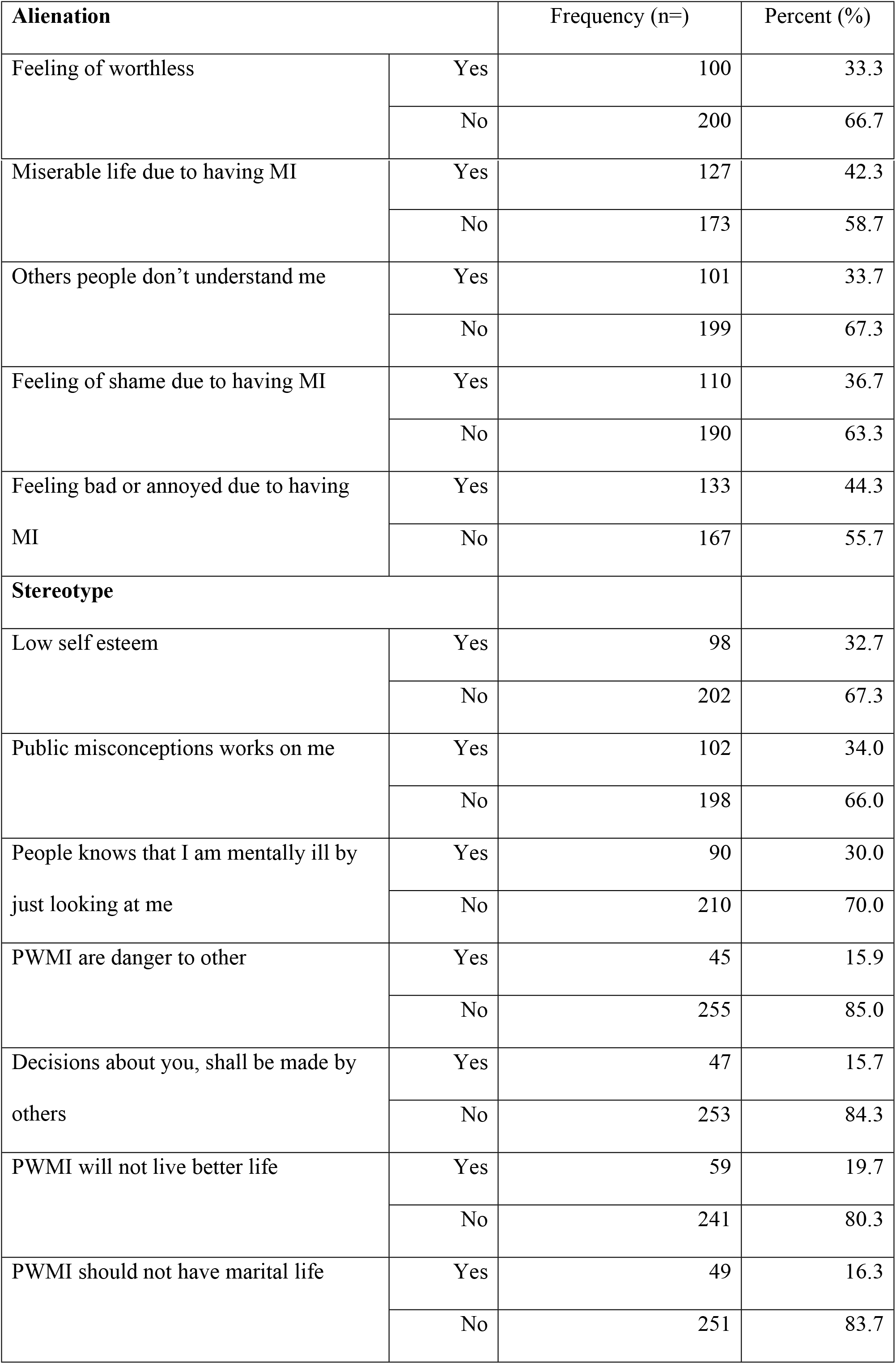
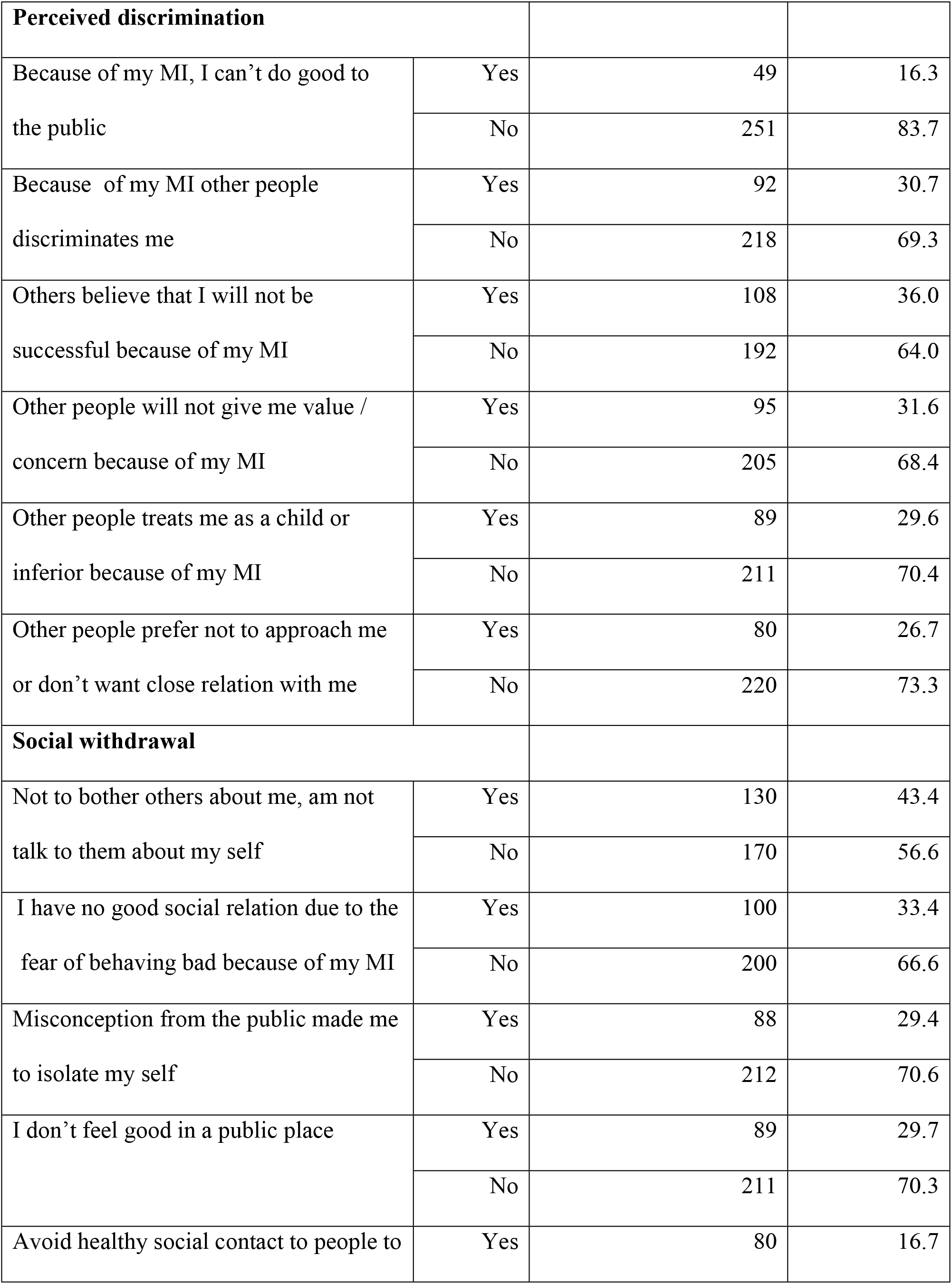
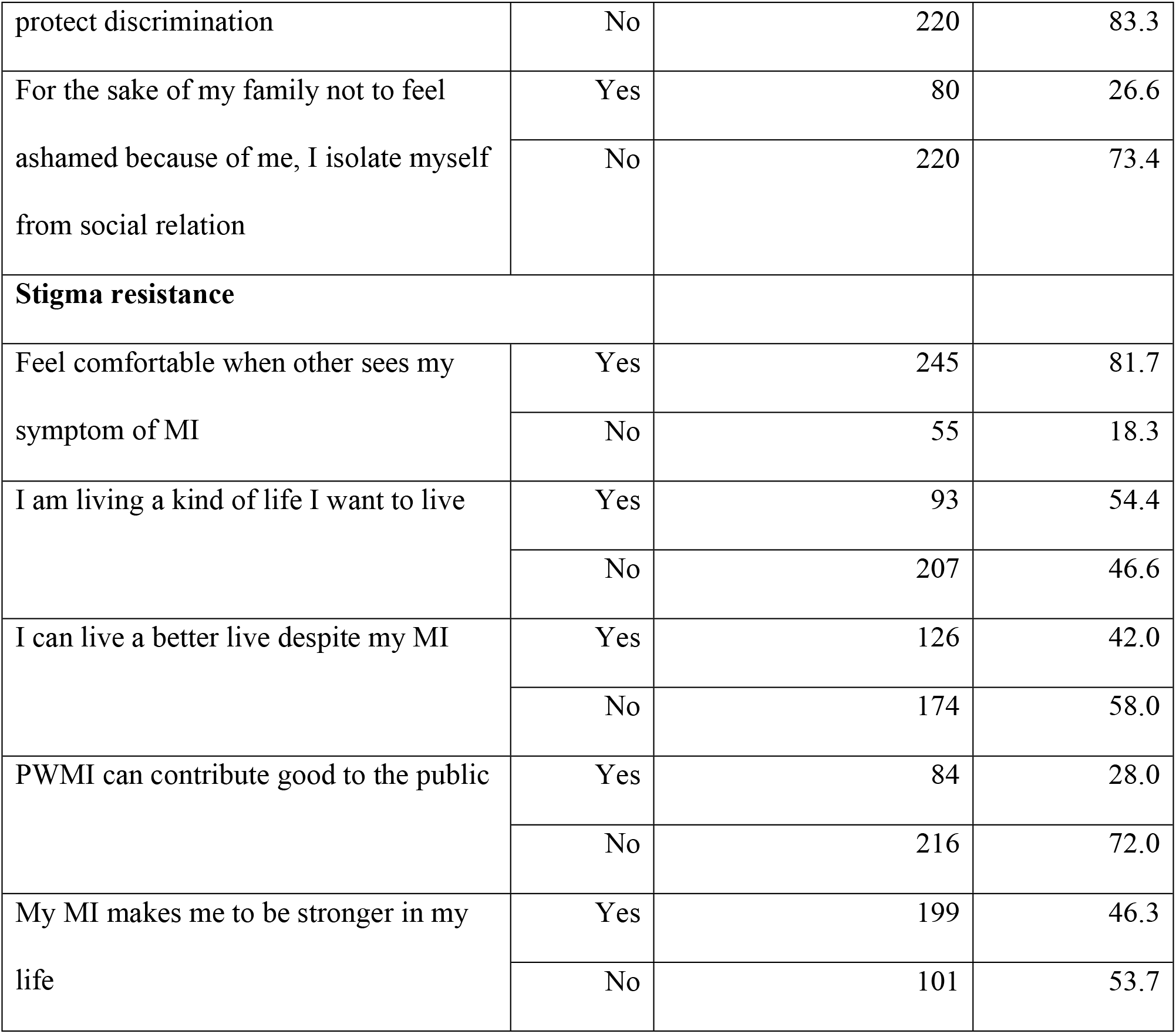
It presents the prevalence of items of overall ISMI score of the patients with mental illness who had history of treatment at psychiatric clinic, 2017.

**Table4.** Morisky scale components with frequency and percentages responded by the patients with mental illness who had history of treatment at psychiatric clinic, 2017.

| Morisky components |  | Frequency<br>(n=) | Percent (%) |
| --- | --- | --- | --- |
| Forgetting their medication | No | 173 | 57.7 |
|  | Yes | 127 | 42.3 |
|  | Total | 300 | 100.0 |
| Careless in taking their medication | No | 194 | 64.7 |
|  | Yes | 106 | 35.3 |
|  | Total | 300 | 100.0 |
| Discontinue their medication when felt better | No | 212 | 70.7 |
|  | Yes | 88 | 29.3 |
|  | Total | 300 | 100.0 |
| Discontinue their medication when felt their illness become worsened | No | 248 | 82.7 |
|  | Yes | 52 | 17.3 |
|  | Total | 300 | 100.0 |

### Bivariate linear regression model

For one by one regression association of each dependent factor with self stigma and discrimination, around 17 variables were associated and candidated for multivariate linear regression. Medication non adherence, having diagnosed with schizophrenia, living with others other than their parents, increased WHODAS score (decreased functional outcome), stopped their treatment during data collection, having poor social support, increased number of admission, having frequent relapse of the illness and staying longer duration without getting treatment for the first time were associated with significantly higher self stigma and discrimination whereas the left variables were associated with lower level of self stigma and discrimination. Table5. It summarizes the univariate factors that were associated with self stigma and perceived discrimination among people with mental illness.

**Table5.**
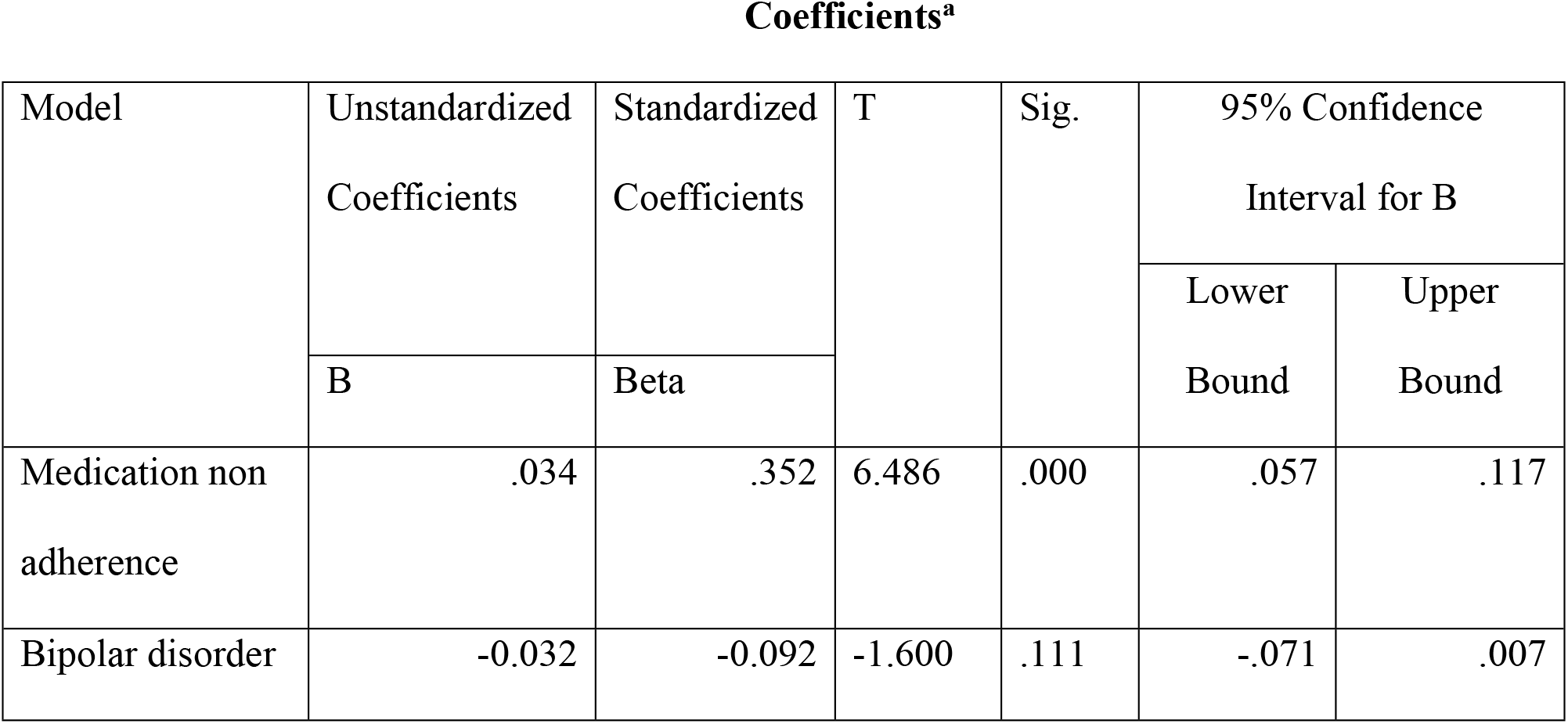
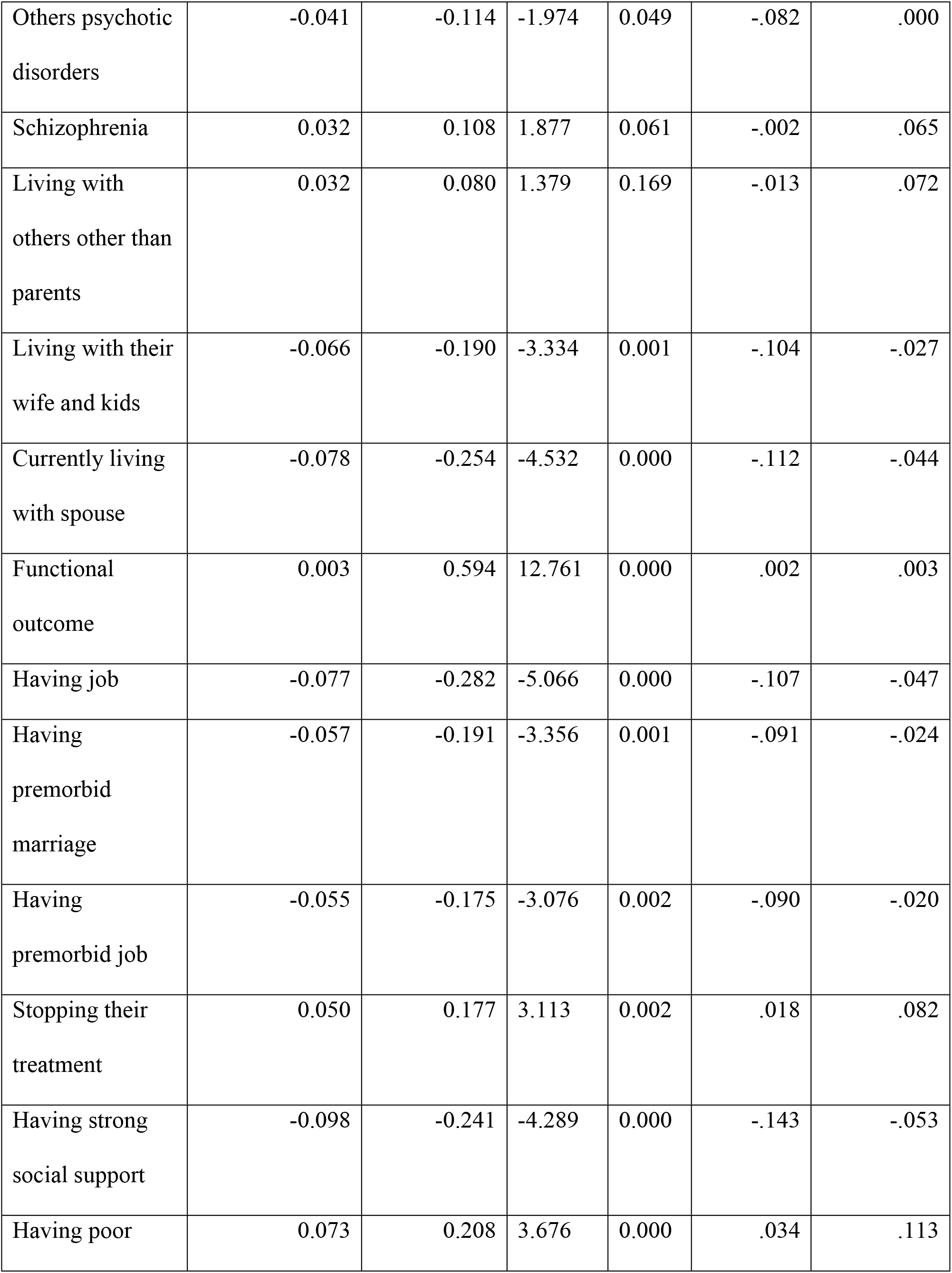
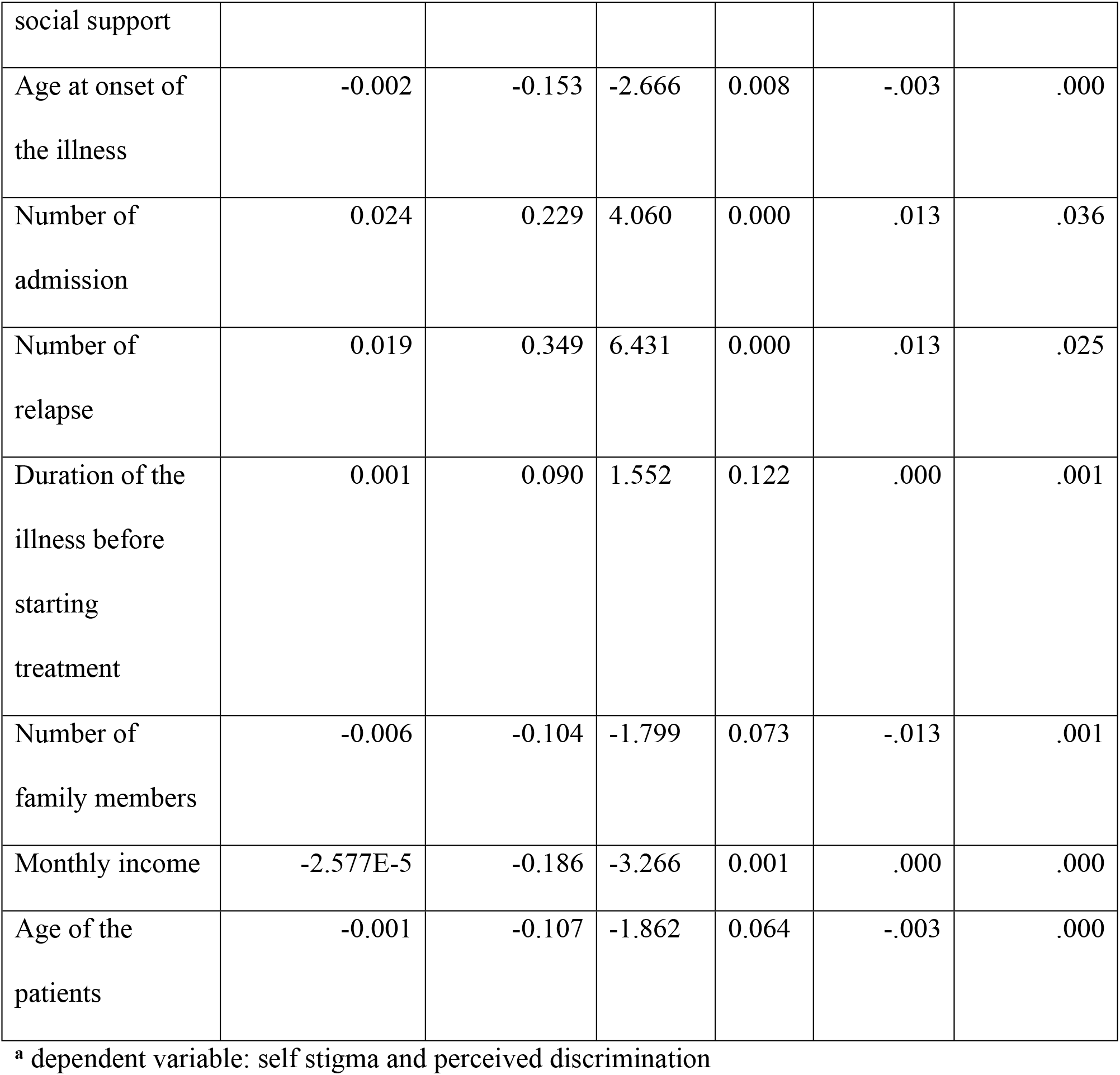
Variables those were associated with self stigma and discrimination in simple regression analysis model of the patients with mental illness who had history of treatment at psychiatric clinic, JUSH, 2016.

### Association of self stigma and level of medication adherence

By dichotomizing level of medication adherence into adherent and non adherent, it was associated with self stigma and perceived discrimination in simple linear regression model and it was candidated to the final adjusted model. In the final model it was significantly associated with self stigma and discrimination. In the Bivariate and multivariate linear regression model, being non adherent to the psychotropic medication was associated with significantly higher self stigma and discrimination, (β=0.034, p<0.0001), (std. β=.084, p<0.05), respectively

**Table6.** It shows the variables that were associated with self stigma and discrimination in the final model of adjusted multivariate linear regression analysis of patients with mental illness who had treatment at psychiatric clinic, JUSH, 2017.

**Coefficients<sup>a</sup>**
|  | Unstandardized<br>Coefficients | Standardized<br>Coefficients | t | Sig. | 95% Confidence Interval for<br>B |  |
| --- | --- | --- | --- | --- | --- | --- |
|  | B | Beta |  |  | Lower<br>Bound | Upper Bound |
| Age of the<br>patient | -.001 | -.091 | -1.899 | .009 | -.002 | .000 |
| Number of<br>relapses | .010 | .183 | 3.868 | .000 | .005 | .015 |
| Increased<br>WHODAS<br>score | .002 | .501 | 10.195 | .000 | .002 | .003 |
| Medication<br>non<br>adherence | .023 | .084 | 1.750 | .021 | .003 | .050 |
| Living with | -.034 | -.099 | -2.081 | .038 | -.067 | -.002 |
| kids and<br>spouse |  |  |  |  |  |  |

<sup>a</sup> dependent variable: self stigma and discrimination

Among adherent respondents, of all level of self stigma, minimal stigma was 63.2% whereas moderate to high stigma was 17.5%. Forty four point one percent of non adherent had moderate to high stigma score. Among the respondents forgetting taking their psychotropic medication, 26.0% had high stigma from all types of stigma whereas the respondents never forget taking of their medication had 5.2% high stigma. From those who were carless in taking their medication in taking their medication according to prescribed, nearly half (47.2%) of the respondents had high self stigma and discrimination. Thirty seven point five (37.5%) of the respondents who had history of medication discontinuation when felt better from their illness without medical advice had moderate to high stigma score. Fifty one point nine percent (51.9%) of the respondents who had history of medication discontinuation when felt their illness worsen without medical advice had moderate to high self stigma score.

As <u>fig.2</u> shows that more than half (53.3%) the respondents scored above the total mean score of self stigma and discrimination are non adherent their prescribed psychotropic medication. <u>Fig.3</u> indicated that though severity of it might be different, self stigma and discrimination was occurred both among adherent and non adherent patients. Accordingly, of all level of self stigma and discrimination minimal stigma was higher (63.4%) among psychotropic medication adherent whereas high self stigma and discrimination was higher (28.0%) among psychotropic medication non adherent than non adherent which was only about 5%.

**Figure2.**
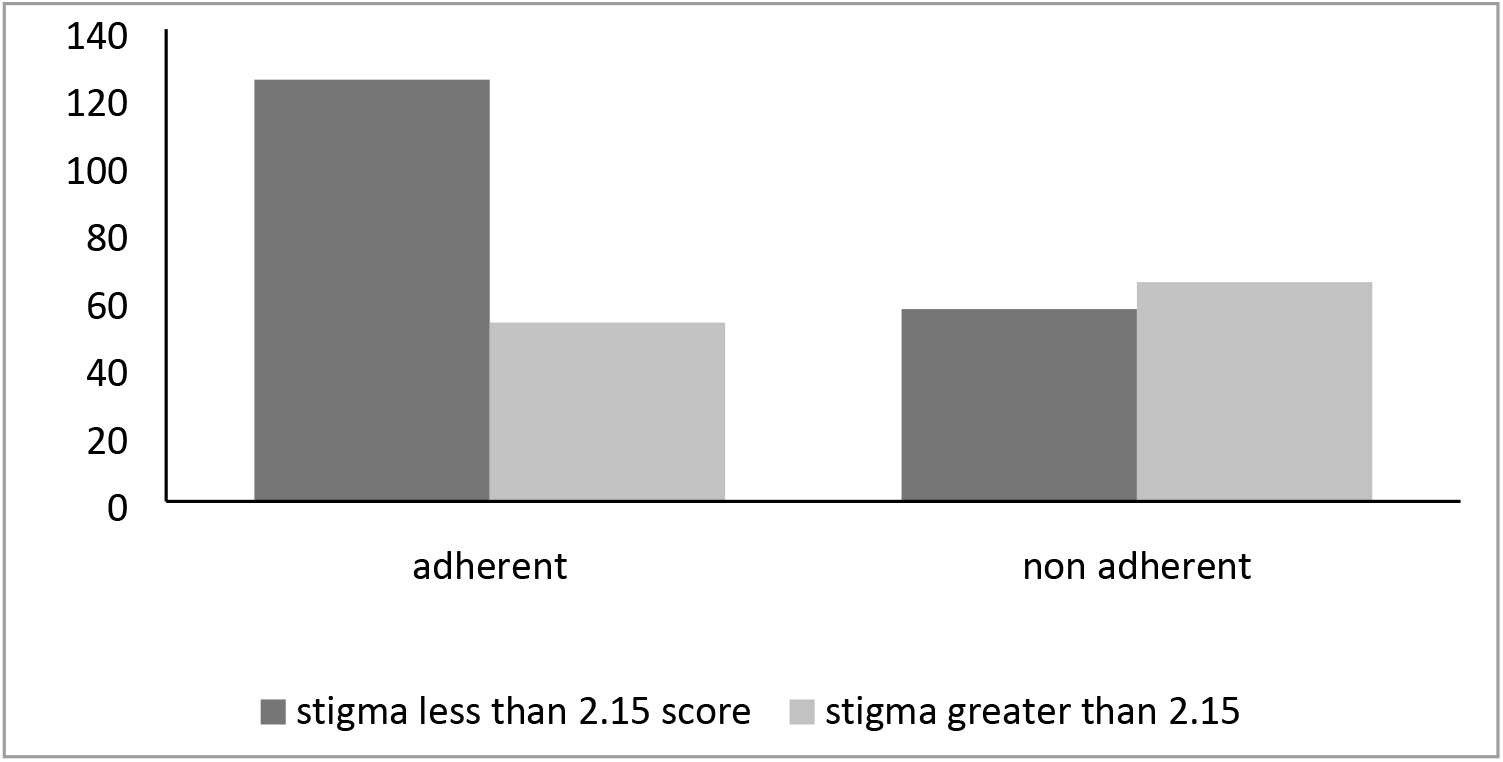
Distribution of the stigma score around the overall mean score of self stigma and discrimination among level of psychotropic medication among patients with mental illness who had treatment at psychiatric clinic, 2017.

**Figure3.**
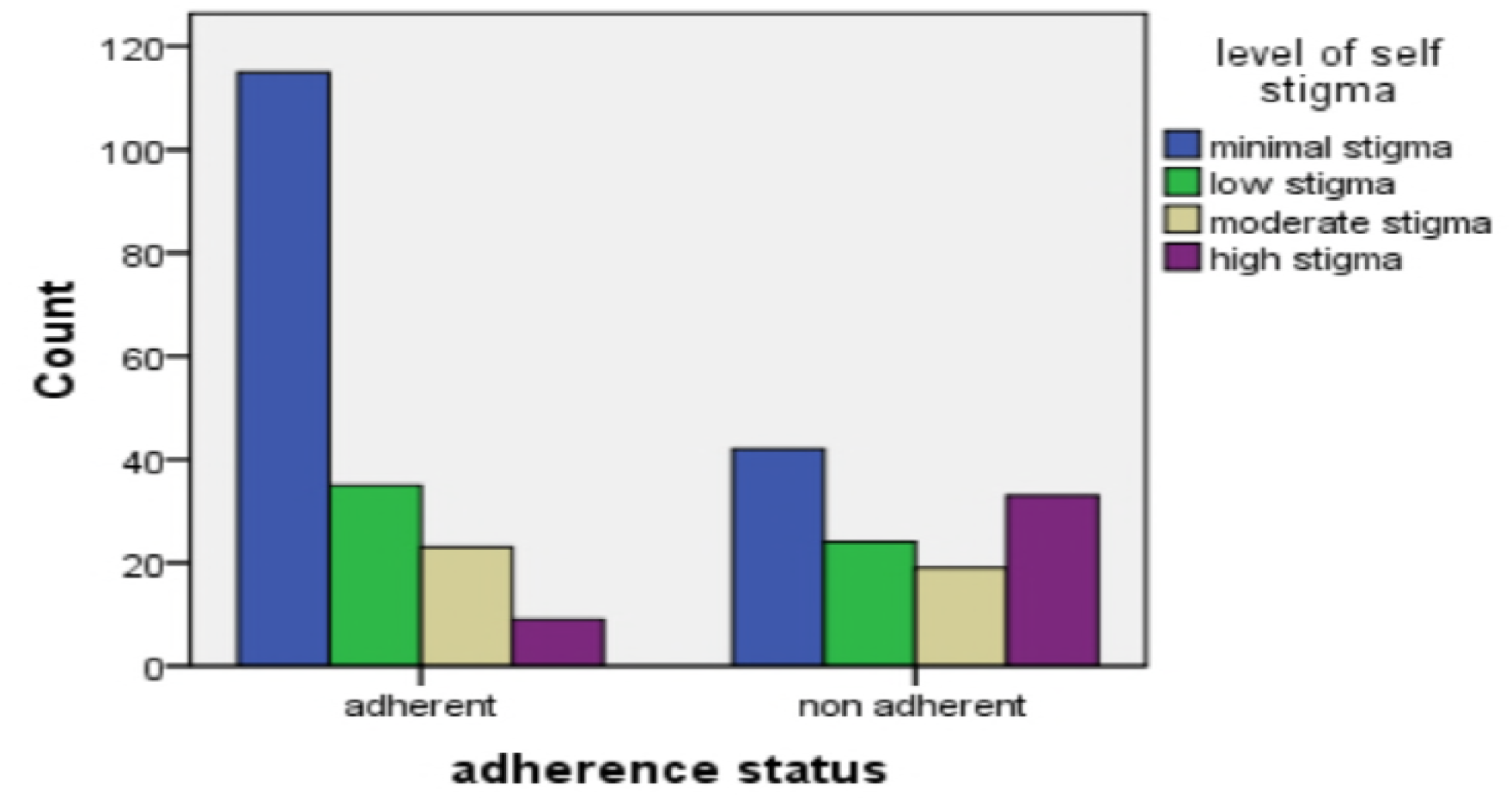
It shows distribution of level of self stigma among adherence status to psychotropic medication among patients with mental illness who had treatment at psychiatric clinic, jimma university teaching hospital, 2017.

When compared with never forgetting of taking the prescribed psychotropic medication, forgetting of taking medication had higher self stigma and discrimination (std. β=.314, p<0.001) and having carelessness in taking of medication had higher self stigma and discrimination (std. β=.364, p<0.001) compared with never be careless. As well, stopping taking of medication when felt well from their illness (std. β=.134, p<0.05) and when felt worse from their illness (std. β=.247, p<0.001) had higher self stigma.

**Figure4.**
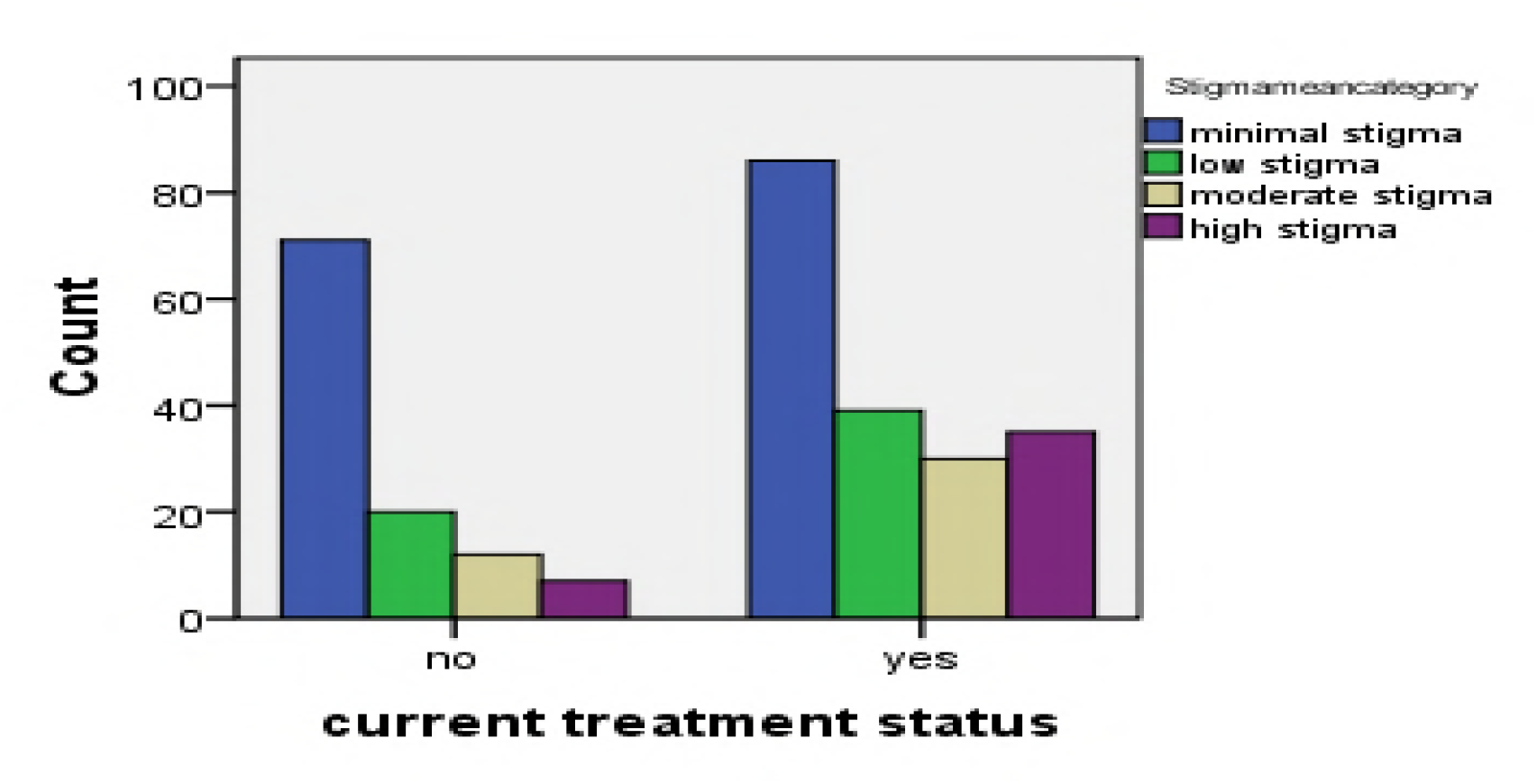
It shows the current treatment status with level of self stigma and perceived discrimination of the patients with mental illness who had history of treatment at psychiatric clinic, JUSH, 2017.

**Table7.** It shows adherence status against self stigma and discrimination of patients with mental illness who had treatment at psychiatric clinic, JUSH, 2017.

|  | Stigma level |  |  |  |  |
| --- | --- | --- | --- | --- | --- |
|  | minimal stigma | low stigma | moderate stigma | high stigma | Total |
| Adherence | 115 | 35 | 23 | 9 | 182 |
|  | 63.2% | 19.2% | 12.6% | 4.9% | 100.0% |
| Non adherence | 42 | 24 | 19 | 33 | 118 |
|  | 35.6% | 20.3% | 16.1% | 28.0% | 100.0% |
| Total | 157 | 59 | 42 | 42 | 300 |
|  | 52.3% | 19.7% | 14.0% | 14.0% | 100.0% |

**Table8.** It shows the frequency and percentage of level of stigma against Morisky medication adherence scale of patients with mental illness who had treatment at psychiatric clinic, JUSH, 2017.

|  | Stigma level |  |  |  | Total |
| --- | --- | --- | --- | --- | --- |
|  | minimal stigma | low stigma | moderate stigma | high stigma |  |
| Forgetting their medication | 43 | 29 | 22 | 33 | 127 |
|  | 33.9% | 22.8% | 17.3% | 26.0% | 100.0% |
| Careless in taking their medication | 34 | 22 | 18 | 32 | 106 |
|  | 32.1% | 20.8% | 17.0% | 30.2% | 100.0% |
| Discontinue their medication when felt better | 38 | 17 | 14 | 19 | 88 |
|  | 43.2% | 19.3% | 15.9% | 21.6% | 100.0% |
| Discontinue their medication when felt their illness become worsened | 16 | 9 | 9 | 18 | 52 |
|  | 30.8% | 17.3% | 17.3% | 34.6% | 100.0% |

In the adjusted regression analysis model, there were others variables which were significantly associated with self stigma and discrimination as of medication non adherence. Those were: age of the patient, number of illness relapse, increased WHODAS score, and living with kids and spouse. As table6 showed, increasing age of the patients and living with kids and spouse was significantly associated with lower level of internalized stigma whereas the rest variables were associated with higher level of internalized stigma.

## Discussion

In this study, the total respondents were 300 with response rate of 98.6% and the majority of the participants were males. Nearly half the respondents were jobless and one third of them followed their education at the level of college and above. The majority of respondents had working diagnosis of schizophrenia followed by major depressive disorder. The mean value of time duration for which the respondents had followed their treatment was 69.11 months and more than half of the respondents had history of discontinuation of their medication without medical advice. Moderate to high self stigma was experienced by around one third of the respondents in this finding and 39.3% participants were non adherent to their prescribed psychotropic medications. Those who had medication adherent behavior had lower self stigma when compared with non adherent.

### Prevalence of self stigma and perceived discrimination

In this finding, the overall self stigma score was 2.15 and around one third (28%) of the respondents had moderate to high self stigma. Near to half (41%) scored greater than overall mean value of ISMI score, which was 2.15. More than one third of the respondents had agreement on feeling of shamefulness due to their mental illness whereas around 43% had low self esteem due to their illness. Equal conclusion was found for the statements of patients with mental illness had difficulty with marital life and mentally ill persons are unable to contribute for the public which was 16.3%. Thirty point seven of the respondents had feeling of socially discriminated due to their illness. The mean score of each of the components of ISMI were alienation, stereotype, discrimination, social withdrawal and stigma resistance: 2.16, 2.11, 2.16, 2.10, and 2.11, respectively.

Comparing with study done in Jimma, lower internalized stigma and discrimination was found in this study whereas higher internalized stigma was found compared with study done in Addis (37, 38). The first study was done among patients with mental illness those only on follow up and its facility based while the current study was done in the community and also took place among all patients who had history of treatment at JUSH without limiting the duration and status of treatment. The second study done in Addis was only among patients with diagnosis of schizophrenia while the current was among patients with all mental illness. Prevalence of self stigma was higher in this study when compared with the studies done in Iran, Europe and Nigeria (15, 36 & 48). Study done in Iran took place among patients with bipolar disorders while study in Nigeria was only among patients with severe mental disorders. This study found that social withdrawal (exclusion from social events) was less experienced than others components of ISMI whereas perceived discrimination and alienation (being inferior) were experienced equally, but the research done in Jimma identified agreement on high alienation followed by stigma resistance (37).

### Prevalence of medication non adherence

In this study, 118 (39.3%) were non adherent to the prescribed psychotropic medication and was close to the study done in JUSH, Jimma, which was 41.2% (46). The study done in India on the prevalence of medication non adherence indicated that around 42% patients were non adherent to their medication which is nearly the same with the current finding (44), but the study done in South Africa showed that higher rates of non adherence which was 63.2%. This difference might be due to study design difference and it also took only among schizophrenic patients (45). The study done in Nigeria indicated that higher prevalence of medication non adherence (48.0%) than the current one and was carried out at facility based and on those patients struggling with illness a sign and symptoms by using self administered questionnaires which might the reasons of difference with the current result (54).

### Association of Self stigma with level of medication adherence

In study significant association was identified between self stigma and perceived discrimination levels of medication adherence. The very noticeable issue in this case was that of level of medication adherence was associated with self stigma in the single linear regression model and also identified as important factor in the final model. Eventhough self stigma and discrimination could occur among all patients, high mean score of self stigma was common among non adherent whereas low mean score was common among adherent than non adherent.

One study done in Sweden to identify the association between experienced stigma and discrimination with medication adherence found that there was no significant association between them in the adjusted model (47), and another study done in Czech Republic, by placing self stigma as independent (predictor factors) for medication adherence, identified that there was negative association between self stigma and medication adherence among schizophrenic patients (41), but the current study assessed association between self stigma medication adherence among patients with mental illness; so that as far as my knowledge, this is the first study on the assessment of association of self stigma and discrimination by taking level of medication adherence as predictor variable among patients with mental illness.

## Conclusion

Higher self stigma and discrimination was found in this study with low mean score of social withdrawal and higher score of alienation and perceived discrimination. There was significant association between overall ISMI score and its subscales and level of medication adherence indicted that becoming non adherent was associated with higher level of self stigma and discrimination. Additionally, all the components of Morisky medication adherence scale were significantly associated with self stigma. Therefore, as one becomes adherent to the prescribed psychotropic medication, level of self stigma and perceived discrimination significantly decreases. Generally, getting and being adherent to mental health treatment and services is not important in reducing and recovery of signs and symptoms of the illness but also important in the recovery of factors which exacerbate or perpetuate the illness, like in this case, self stigma and discrimination associated with misconception about mental illness.

### Limitation of the study

To get the study participants, we used general survey method on patients with mental illness who had history of treatment at psychiatric clinic which might lead to recall bias and great variability in information of the current data (can significantly affect the reliability and validity of the result). In this study, we assessed only internalized stigma and perceived discrimination but also it would be better if others stigma and discrimination (social and institutional stigma and applied discrimination) additionally addressed to identify its magnitude and association with level of medication adherence among patients with mental illness. We assessed the association of self stigma and level of medication adherence cross sectionally but really to identify the impact of treatment (medication adherence) on the level of self stigma and perceived discrimination, using others study design than currently used would be better. Eventhough study’s aim was to address and assess status of all the patients with mental illness who had history of treatment at psychiatric clinic, JUSH; we were unable to address those patients who left the town for different reasons. This study excluded all the patients who got treatment from the hospital for less than six months duration, but to assess level of self stigma with impact of treatment, it would be better if they were included.

## Acknowledgement

My deepest gratitude and thanks goes to my advisors for their heartfelt advising, commenting and showing direction by giving motivation for me to go ahead. I would like to extend my thanks to health extension worker of the town, kebele administration stakeholders and all bodies that were with me in different activities to do this study.

